# Age, sex, and puberty related development of the corpus callosum: a multi-technique diffusion MRI study

**DOI:** 10.1101/187724

**Authors:** Sila Genc, Charles B Malpas, Gareth Ball, Timothy J Silk, Marc L Seal

**Author notes:** equal senior author (Timothy J Silk, Marc L Seal). Corresponding author: Sila Genc Department of Paediatrics, The University of Melbourne, Parkville VIC 3052, Australia.; E; ORCID ID: 0000-0002-4624-7071. Secondary corresponding author: Tim Silk, E. **Abbreviations** AFD: Apparent Fibre Density AIC: Akaike Information Criterion BMI: Body mass index DTI: Diffusion tensor imaging DWI: Diffusion-weighted imaging CI: Confidence interval CMIND: Cincinnati MR Imaging of NeuroDevelopment CSD: Constrained Spherical Deconvolution FA: Fractional anisotropy FBA: Fixel-based analysis FOD: Fibre orientation distribution GLM: General linear model MRI: Magnetic resonance imaging MD: Mean Diffusivity NICAP: Neuroimaging of the Children’s Attention Project NODDI: Neurite Orientation Density Dispersion Imaging ODI: Orientation Dispersion Index PDS: Pubertal development scale TE: Echo-time TR: Repetition time *v_ic_*: Intra-cellular volume fraction.

## Abstract

**Purpose:** The corpus callosum is integral to the central nervous system, and continually develops with age by virtue of increasing axon diameter and ongoing myelination. Magnetic resonance imaging (MRI) techniques offer a means to disentangle these two aspects of white matter development. We investigate the profile of microstructural metrics across the corpus callosum, and assess the impact of age, sex and pubertal development on these processes.

**Methods:** This study made use of two independent paediatric populations. Multi-shell diffusion MRI data were analysed to produce a suite of diffusion tensor imaging (DTI), neurite orientation density and dispersion imaging (NODDI), and apparent fibre density (AFD) metrics. A multivariate profile analysis was performed for each diffusion metric across 10 subdivisions of the corpus callosum.

**Results:** All diffusion metrics significantly varied across the length of the corpus callosum. AFD exhibited a strong relationship with age across the corpus callosum (partial *η*^2^ = .65), particularly in the posterior body of the corpus callosum (partial *η*^2^ = .72). In addition, females had significantly higher AFD compared with males, most markedly in the anterior splenium (partial *η*^2^ = .14) and posterior genu (partial *η*^2^ = .13). Age-matched pubertal group differences were localised to the splenium.

**Conclusion:** We present evidence of a strong relationship between apparent fibre density and age, sex, and puberty during development. These results are consistent with *ex vivo* studies of fibre morphology, providing insights into the dynamics of axonal development in childhood and adolescence using diffusion MRI.

**Target journals:** Brain Structure & Function; HBM; NeuroImage; Developmental Cognitive Neuroscience

## 1. Introduction

Brain development throughout childhood and adolescence is a dynamic process, with maturation of the *white matter* characterised by ongoing axonal development (LaMantia and Rakic 1990) and myelination (Yakovlev and Lecours 1967). These two integral components of the white matter, the axon and myelin sheath, contribute to the microstructural organisation of a white matter fibre bundle and should be assessed separately (Paus 2010). However, it has not been possible to assess, *in vivo*, the relative contribution of these features until the recent introduction of sophisticated diffusion-weighted imaging (DWI) acquisition and modelling techniques.

The corpus callosum is an integral part of the central nervous system, providing a physical pathway for information transfer between the two hemispheres of the brain. The corpus callosum is a highly organised bundle of fibres that converges at the midline, and projects out to the cortical surfaces. Axon count in the corpus callosum stabilises in the early post-natal period (LaMantia and Rakic 1990), quickly reaching adult levels and serving as a structural scaffold for further neuronal maturation. From this point, evidence suggests that continuing axon diameter growth and myelination is synchronised (Sherman and Brophy 2005; Nave 2010). A recent lifespan assessment of the g-ratio (the ratio of the inner axonal diameter to the total outer diameter) in the corpus callosum revealed a stable trajectory with age, further evidencing this coupled growth (Berman et al. 2017).

Fibre composition varies across the corpus callosum, with differing proportions of small and large diameter fibres across callosal subregions (Aboitiz et al. 1992). The posterior (splenium) and anterior (genu) segments are composed of a high density of small diameter fibres, whereas the body has a higher proportion of large diameter fibres. Differences in myelin composition are also apparent across the corpus callosum, as magnetic resonance imaging (MRI) derived estimates of myelin volume fraction are highest in the genu and lowest in the body (Bjornholm et al. 2017; Stikov et al. 2015).

*In vivo* estimates of white matter organisation are most commonly derived from diffusion tensor imaging (DTI). Most commonly, fractional anisotropy (FA) is used to estimate the degree of anisotropic water diffusion along a fibre bundle, where high FA can signify greater organisation of white matter fibre tracts. In studies of development, DTI applications have revealed that the white matter becomes more highly organised with age (Lebel and Beaulieu 2011), and that age-related development of the corpus callosum follows a known developmental course (Lebel et al. 2012; Lebel et al. 2008). Whilst useful as a sensitive marker of microstructural alterations, DTI metrics are not specific enough to disentangle the physical properties of a fibre bundle. Crossing fibres, glial abundance/loss, partial voluming, and orientation dispersion can conflate the inference of axon and myelin ‘organisation’ (Jones et al. 2013; Beaulieu 2002).

Recent advancements in modelling and acquisition methods offer to disentangle multiple fibre properties using clinically feasible acquisitions. Neurite orientation dispersion and density imaging (NODDI) (Zhang et al. 2012) is a compartment modelling technique that estimates neurite density as the intra-cellular volume fraction (*v*_*ic*_), as well as the bending and fanning of axons as the orientation dispersion index (ODI). Apparent fibre density (AFD) is an alternative metric that quantifies the total intra-axonal volume fraction, to estimate axon density per unit of tissue (Raffelt et al. 2017; Raffelt et al. 2012). Higher AFD in a given fibre bundle can reflect larger axon diameter, or higher local axon count. Using such metrics instead of DTI-based measures allow a more specific interrogation of axonal properties, which are useful as axonal conduction velocity is determined by axon diameter, whereby larger axons conduct faster (Horowitz et al. 2015). Estimating axon diameter, however, using current clinical MRI systems is not feasible for small axons, which make up a large proportion of the brain (Nilsson et al. 2017). Therefore the quantification of white matter fibre density estimates are warranted.

We focus our efforts on examining corpus callosum development using several commonly derived and recently introduced diffusion MRI metrics. We first provide a detailed examination of how several diffusion metrics vary across the length of the corpus callosum, with comparison to previously reported profiles in adults. Subsequently, we examine how these relationships vary as a function of age, sex and pubertal stage during development. We utilise two independent developmental samples; (1) a developmental population (aged 4.1 - 18.9 years), and (2) an age-matched group of children at the cusp of pubertal onset (aged 9.6 - 11.9 years).

## 2. Methods

### Cohorts

Two independent, typically-developing cohorts of children and adolescents were used for the current study:

The Cincinnati MR Imaging of NeuroDevelopment (CMIND) study comprised a sample of children from early infancy to late adolescence, based in Cincinnati, USA. Data acquired at a single site, the Cincinnati Children∙s Hospital Medical Center (CCHMC), were accessed from their online data repository (Holland SK 2015). Full description of the recruitment process and MRI scanning protocol is detailed online (https://cmind.research.cchmc.org/). Only participants between the ages of 4 - 19 with DWI data were included for analysis in this study (N = 85, M = 10.80 years, 45 female).

The Neuroimaging of the Children’s Attention Project (NICAP) study comprised a community sample of children aged 9 - 12 years, recruited from 43 socio-economically diverse primary schools distributed across the Melbourne metropolitan area, Victoria, Australia. A detailed protocol has been described previously (Silk et al. 2016). This narrow age band of children included typically developing pre-pubertal (N = 44, *M* = 10.41 years, 13 female) and pubertal children (N = 30, *M* = 10.45 years, 18 female), as defined by the Pubertal Development Scale (PDS). For further details on pubertal group classification, see (Genc et al. 2017b). Only participants that had both DWI data and PDS data were included in the current study (N = 74, *M* = 10.40 years, 31 female). This study was approved by The Royal Children’s Hospital Melbourne Human Research Ethics Committee (HREC #34071).

### Image Acquisition

CMIND children underwent MRI at 3.0T on a Phillips Achieva TX. In brief, diffusion-weighted images were obtained with a spatial resolution of 2.0 x 2.0 x 2.0 mm, acquisition matrix = 112 x 109, bandwidth = 1753 Hz, 60 slices, along 61 directions. Two DWI shells were acquired, one b = 1000 s/mm^2^ shell (relaxation time [TR] = 6614 ms, echo-time [TE] = 81 ms) and one b = 3000 s/mm^2^ shell (TR = 8112 ms, TE = 104 ms). The total acquisition time for both shells was 21m 27s.

NICAP children underwent MRI at 3.0T on a Siemens Tim Trio at The Melbourne Children’s Campus, Melbourne, Australia. Briefly, three diffusion-weighted shells were acquired: (1) b = 2800 s/mm^2^, 60 directions; (2) b = 2000 s/mm^2^, 45 directions; and (3) b = 1000 s/mm^2^, 25 directions. All shells had a voxel size of 2.4 x 2.4 x 2.4 mm, acquisition matrix = 110 x 100, bandwidth = 1758 Hz, TR = 3200 ms, TE = 110 ms. A multi-band acceleration factor of 3 significantly reduced the acquisition time (9m 23s). An additional reverse phase-encoded image was collected to assist with susceptibility induced distortion correction (total acquisition time = 10m 33s).

### Data pre-processing and analysis

For CMIND, a total of 72 participants had sufficient data for DTI and NODDI analyses (*M* = 10.42, *SD* = 3.99, 36 female). Images were pre-processed and a tensor-based population-specific template (Zhang et al. 2006) was generated as previously described in Genc et al. (2017a). Subsequently, fractional anisotropy (FA) and mean diffusivity (MD) maps were generated in template space from the b = 1000 s/mm^2^ data using *TVtool*. NODDI maps (*v*_*ic*_ and ODI) were generated from the combined low and high b-value data (b = 1000, 3000 s/mm^2^) using the NODDI toolbox (Zhang et al. 2012), and then transformed to template space using DTI-TK (Zhang et al. 2006). For the AFD analysis, a total of 85 participants with high b-value data (b = 3000 s/mm^2^) were included for analysis in this study (*M* = 10.80, *SD* = 3.90, 45 female). Data were processed using MRtrix3 (https://github.com/mrtrix3). Data were denoised, corrected for distortions due to eddy currents and bias fields, normalised for variations in global intensity, and upsampled by a factor of 2. A fibre orientation distribution (FOD) map was generated for each participant, and subsequently a study specific FOD template was generated using one male and one female for each age between 5 - 18 years (total of 28 participants). Each participant’s FOD map was transformed to template space, and the first volume of the FOD map was extracted to estimate apparent fibre density (AFD) in the corpus callosum.

For NICAP, a total of 74 participants had sufficient DWI data for further analyses (*M* = 10.40 years, 31 female). A population-specific template was generated using the low b-value data (as above). FA and MD maps were generated from the b = 1000 s/mm^2^ data using *TVtool*. Using the combined b = 1000 and b = 2000 s/mm^2^ data, *v*_*ic*_ and ODI maps were generated using the NODDI toolbox and transformed to template space using DTI-TK. Following visual inspection, a total of 68 *v*_*ic*_ maps and 71 ODI maps were included for further analysis. Processing of the higher b-value data (b = 2800 s/mm^2^) for AFD estimation was performed on 74 children as described previously in (Genc et al. 2017b) and as above.

In summary, data used for the CMIND cohort comprised: FA, MD, *v*_*ic*_ and ODI data (N = 72, *M* = 10.42 years, 36 female), and AFD data (N = 85, *M* = 10.80, 45 female). Data used for the NICAP cohort comprised: FA, MD, and AFD data (N = 74, *M* = 10.40 years, 31 female), *v*_*ic*_ data (N = 68, M = 10.40 years, 30 female) and ODI data (N = 71, M = 10.42 years, 31 female).

### Corpus callosum parcellation

The corpus callosum segmentation was performed according to Aboitiz et al. (1992), whereby the mid-sagittal slice of the corpus callosum was divided along its length to generate 10 segments: genu (G1, G2, G3), body (B1, B2, B3), isthmus (ISTH) and splenium (S1, S2, S3). This is represented as a colour-coded map in *Fig 1*, and tract projections were also generated purely for visualisation (*Fig 3b*). Regions of interest (ROIs) were drawn on the population-based tensor template for DTI and NODDI quantification, and on the FOD template for AFD quantification, for each respective cohort using *mrview*. A mean value per segment for each diffusion metric was calculated for statistical analysis.

**Fig. 1.**
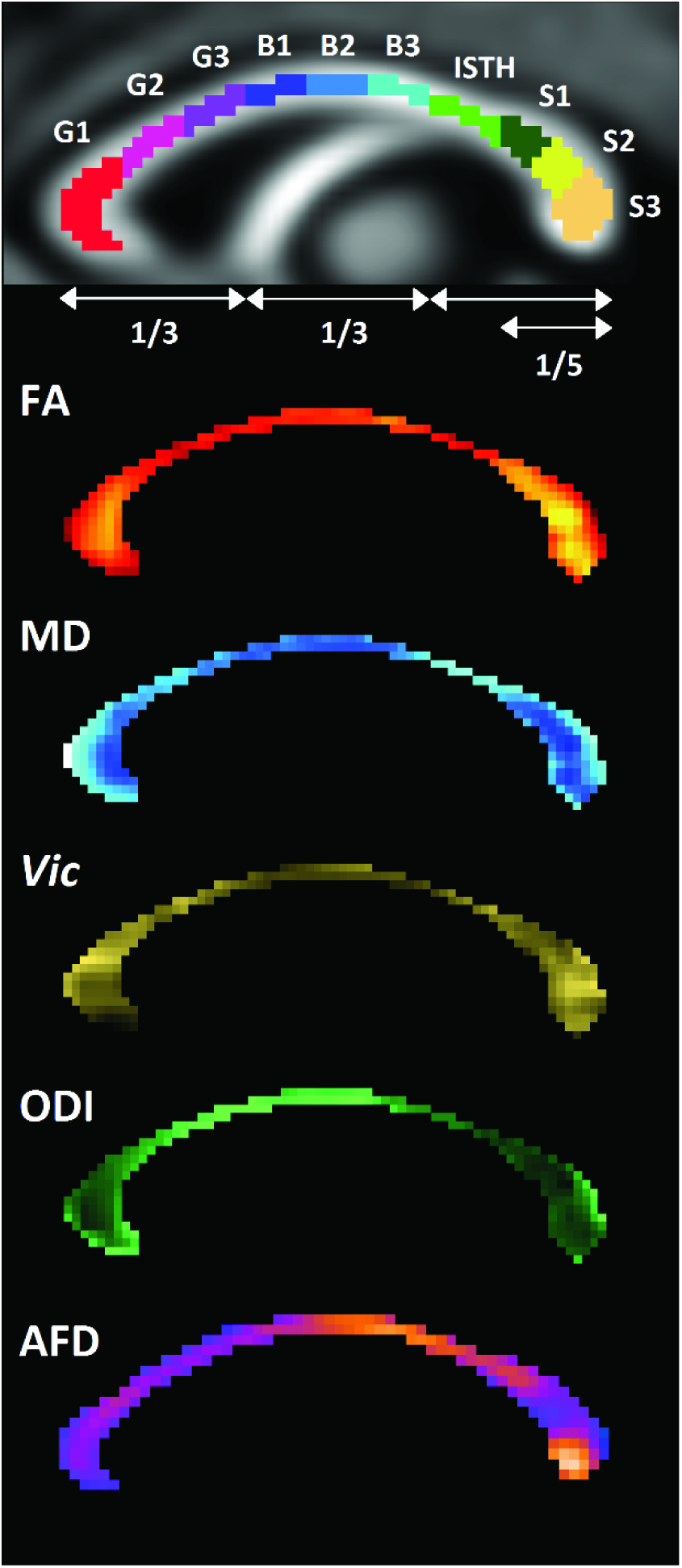
Corpus callosum parcellation on mid-sagittal slice of FOD template, alongside heat-maps of diffusion metric variability across the corpus callosum. Brighter colours correspond to higher values

### Statistical analyses

Corpus callosum profiles were examined using profile analysis via multivariate analysis of covariance (MANCOVA) (SPSS, version 24). Corpus callosum segment was entered as the within-subjects factor for each analysis. Age and sex were included as between-subjects covariates. Main effects and interactions are reported for CMIND and NICAP cohorts.

Statistically significant interactions involving segment were further investigated using univariate linear models separately for each segment (R, version 3.4.1). For age relationships, in addition to first order linear models, second and third order models were also computed and Akaike Information Criterion (AIC) values were reported. The coefficient of determination (adjusted *R^2^*) was computed to understand the proportion of variance explained by the model. Partial eta-squared (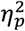) is reported as an estimate of effect size.

Multivariate profile analysis was also performed for the pubertal group comparisons in the NICAP cohort, as information on pubertal status derived from the PDS was available for this cohort. Sex and pubertal group were included as between-subjects factors. Follow-up univariate linear models were computed for each segment, to examine the relationship between pubertal group and AFD. In addition, we performed an exploratory group comparison to assess step-wise changes in AFD after generating a new variable, pubertal-type, which consisted: pre-pubertal male; pubertal male, pre-pubertal female, and pubertal female.

All graphical visualisations were carried out in R (version 3.4.1). Coloured diffusion metric profiles were plotted as 95% confidence intervals (CIs) and adjusted for repeated measures across the 10 corpus callosum segments.

We made use of a strict statistical threshold for significance, due to multiple comparisons between groups and corpus callosum segments. As a result, we have defined our threshold of statistical significance as *p* < .005.

## 3. Results

### Corpus callosum profiles

For both CMIND and NICAP datasets, the variability of diffusion metrics across the corpus callosum are visualised as heat-maps (*Fig 1*), and as profiles plotted across the ten segments of the corpus callosum (*Fig 2a*). For the CMIND cohort, age-related development of the profiles are visualised by grouping data into three age groups (4.1 - 8.1; 8.3 - 12.3; 12.3 - 18.8 years).

**Fig. 2.**
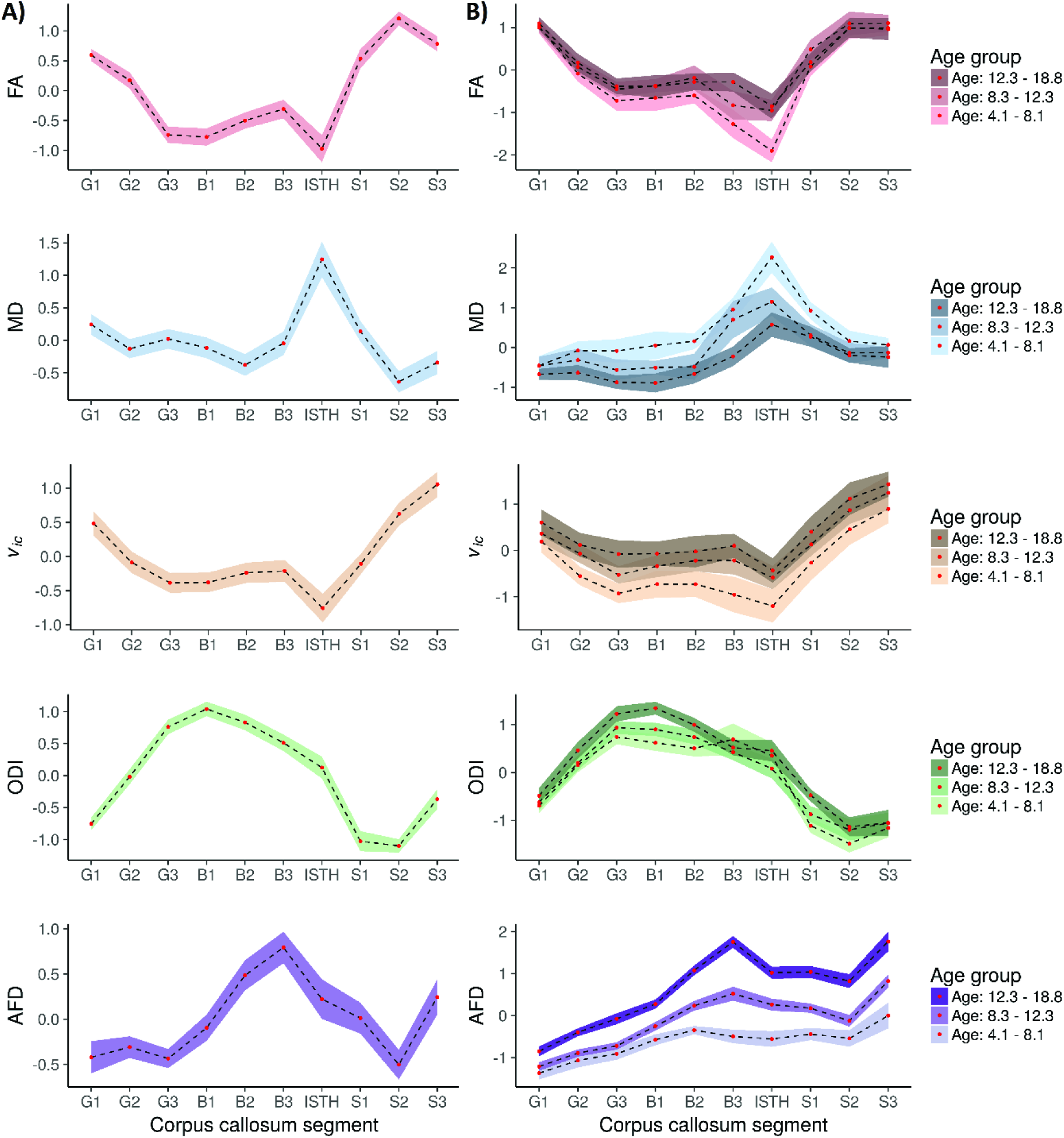
Profiles of diffusion metrics across the 10 segments of corpus callosum in A) NICAP, and B) CMIND cohorts. CMIND groups are split into 3 age groups for visualisation, where the coloured ribbon is darker with older age group. Coloured ribbons represent 95% CIs

In the CMIND cohort, a high-low-high profile was observed for both FA and *v*_*ic*_ across the 10 segments, with profiles generally translating upwards suggesting an increase in FA and *v*_*ic*_ with age. MD followed a low-high-low profile, peaking in the isthmus. ODI peaked at B1, and followed a downward pattern reaching a minima at the splenium. AFD showed most marked differences between age-groups, with an anterior-posterior increase peaking at B3.

Multivariate profile analysis for diffusion metrics in the CMIND cohort confirmed the profiles listed above. These results are reported in *Table S1*. In general, all metrics had a statistically significant main effect for segment, indicating variability across the segments. Main effect of age was significant for MD, *F*(1,69) = 29.1, *p* < .001, 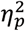 = .30, ODI, *F*(1,69) = 21.5, *p* < .001, 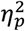 = .76, and AFD, *F*(1,82) = 154.14, *p* < .001, 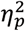 = .65. The segment by age interaction effect was statistically significant for ODI, *F*(9,61) = 4.94, *p* < .001, 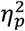 = 42, and AFD, *F*(9,74) = 21.55, *p* < .001, 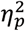 = .72, indicating that the profiles were not parallel across different ages.

The diffusion metric profiles over the corpus callosum in NICAP were similar to that of CMIND. We observed that FA and *v*_*ic*_ both exhibited a high-low-high profile across the 10 segments, with a minimum in the isthmus and maximum in the splenium. MD exhibited a low-high-low profile, peaking in the isthmus. We observed a smoother profile of low-high-low ODI from the genu to the isthmus, before reaching a minima at S2 and rising again at S3. AFD had a low-high-low profile from the genu to S2, peaking at B3 and rising again at S3. Multivariate profile analysis for diffusion metrics in the NICAP cohort are reported in *Table S2*. There was no evidence for a significant main effect of age, sex, or interactions between segment and age, and segment and sex, for any of the diffusion metrics.

Given that AFD had the greatest magnitude of main effect of age in the CMIND cohort, *F*(1,82) = 154.14, *p* < .001, 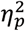 = .65, and interaction with age, *F*(9,74) = 21.55, *p* < .001, 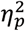 = .72, we subjected each corpus callosum segment to further analysis (below).

### Age relationships

Linear models were computed to further investigate the relationship between AFD and age in each corpus callosum segment separately. For each segment, AFD was regressed onto age (with sex included as a covariate). As shown in *Table 1*, AFD in the body of the corpus callosum had the strongest relationship with age (B2: 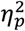 = .60 and B3: 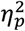 = .72). In addition to first order (linear) effects, we also modelled second order (quadratic) and third order (cubic) effects. Overall the quadratic and cubic models did not result in a better fit as indicated by both AIC and adjusted *R^2^* values (*Table S4*), therefore only the linear relationships are reported and visualised in *Fig 3a*.

**Table 1.** Relationship between AFD with age and sex, across the 10 segments of the corpus callosum

| Region | Overall | Age |  |  | Sex |  |  |
| --- | --- | --- | --- | --- | --- | --- | --- |
| | *Adjusted R<sup>2</sup>* | *F(1,82)* | *p* | $\eta_p^2$ | *F(1,82)* | *p* | $\eta_p^2$ |
| G1 | .24 | 20.6 | < **.001** | .22 | 7.2 | .009 | .08 |
| G2 | .38 | 42.9 | < **.001** | .36 | 10.3 | **.002** | .11 |
| G3 | .47 | 63.3 | < **.001** | .45 | 12.3 | < **.001** | .13 |
| B1 | .42 | 55.7 | < **.001** | .42 | 7.5 | .008 | .08 |
| B2 | .60 | 118.9 | < **.001** | .60 | 6.6 | .01 | .07 |
| B3 | .71 | 205.5 | < **.001** | .72 | 5.6 | .02 | .06 |
| ISTH | .54 | 91.0 | < **.001** | .54 | 8.1 | .006 | .09 |
| S1 | .57 | 98.5 | < **.001** | .56 | 13.2 | < **.001** | .14 |
| S2 | .49 | 72.4 | < **.001** | .48 | 9.0 | **.004** | .10 |
| S3 | .51 | 80.1 | < **.001** | .51 | 10.9 | **.001** | .12 |

Adjusted $R^2$ values were calculated using a linear model. F-statistic and p-value for age and sex reported from
ANOVA, where bold values indicate $p < .005$ . Partial eta squared ( $\eta_p^2$ ) reported as an estimate of effect size.

**Fig. 3.**
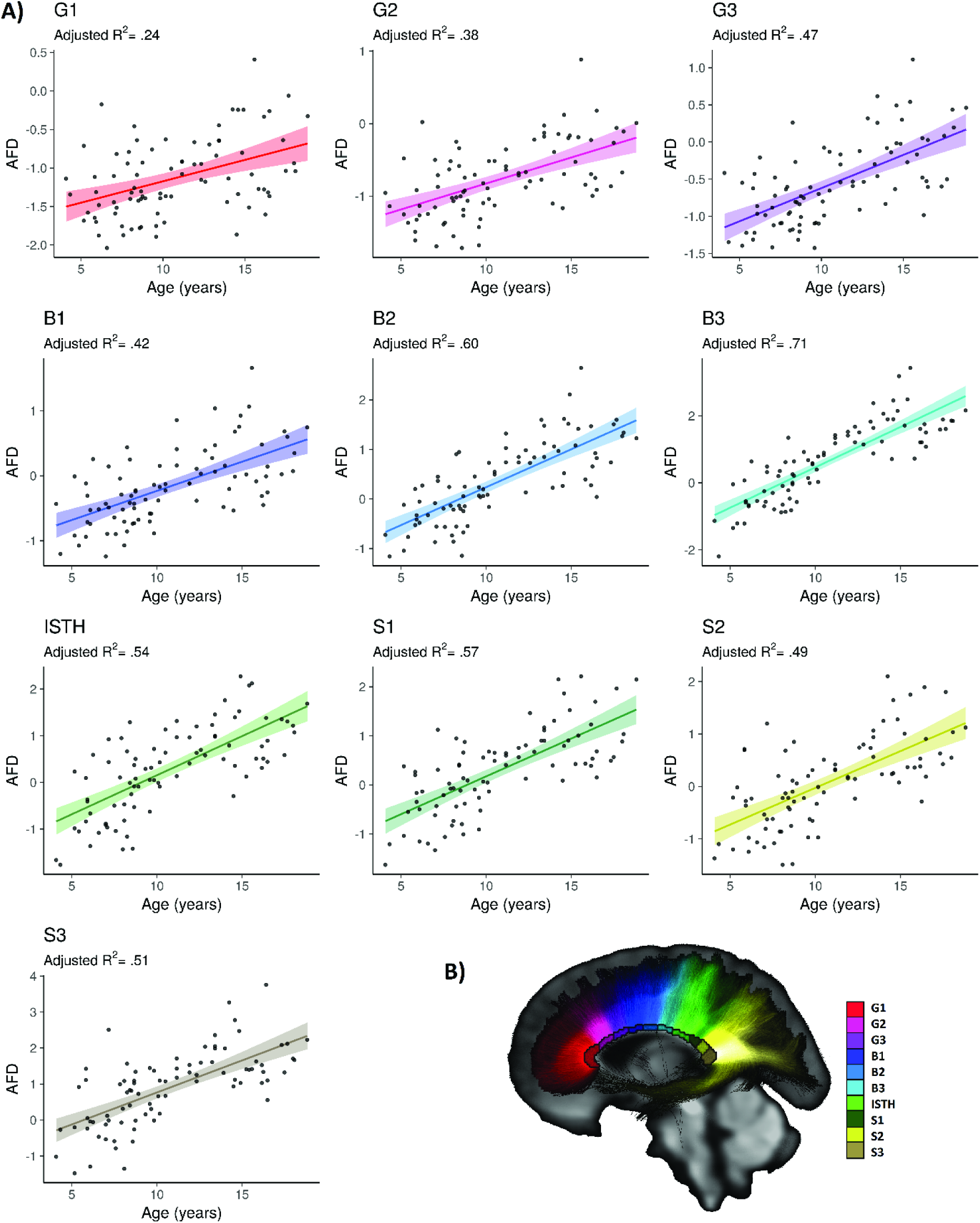
Relationship between AFD and age across the corpus callosum A) Linear relationship between AFD and age, where adjusted *R^2^* values are adjusted for sex, and shaded areas around the line of fit represent 95% CIs. B) Anatomical position and tract projections shown for each corpus callosum segment

### Sex differences

For the CMIND cohort, there was a statistically significant main effect of sex across the callosal segments, *F*(1,82) = 15.62, *p* < .001, 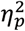 = .16, suggesting that females had higher AFD than males (*Fig S1a*). The segment by sex interaction, however, was not statistically significant for AFD, *F*(9,74) = 0.61, *p* = .78, 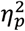 = .07.

Overall, age had a stronger effect on AFD across the 10 segments (.22 < 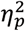 < .72), compared to the relatively weaker effect of sex (.06 < 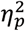 < .14). The strongest evidence for sex differences was observed in G3 (*p* < .001, 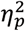 = .13), and S1 (*p* < .001, 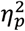 = .14) (*Table 1*).

### Influence of puberty

We performed an investigation into the influence of pubertal group on AFD across the corpus callosum in the NICAP cohort. Multivariate profile analysis revealed a significant segment by pubertal group interaction for AFD, *F*(9, 62) = 3.12, *p* = .004, 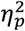 = .31. Evidence of pubertal group differences was only apparent in S2, *F*(1,70) = 10.3, *p* = .002, 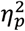 = .13, and S3, *F*(1,70) = 8.98, *p* = .004, 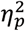 = .11.

In addition, we performed an exploratory group comparison to assess differences in AFD across pre-pubertal and pubertal males and females (pubertal-type). There was evidence for a segment by pubertal-type interaction, *F*(27,192) = 1.91, *p* = .007, 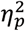 = .21. The main effects of pubertal-type were statistically significant in the regions S2, *F*(3, 70) = 5.1, *p* = .003, 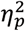 = .18, and S3, F(3, 70) = 6.45, *p* = .001, 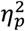 = .22 (*Fig S1b*), suggesting a step-wise progression of AFD.

## 4. Discussion

The current study presents a comprehensive investigation of the influence of age, sex, and puberty on corpus callosum development using a number of DWI modelling and analysis techniques in two independent developmental samples.

### Corpus callosum profiles

The profiles across the corpus callosum for FA and MD are consistent with previous reports in adults (Caminiti et al. 2013; Bjornholm et al. 2017). The profile of *v*_*ic*_ followed a very similar pattern to FA, although *v*_*ic*_ did not show an interaction with age as each segment increased similarly with age. Since the corpus callosum is a coherently organised bundle of white matter fibres, it is possible that *v*_*ic*_ does not substantially improve the characterisation of neurite properties compared with FA (Genc et al. 2017a).

Upon visual inspection of the FA and *v*_*ic*_ profiles alone might lead to the interpretation that the genu and splenium have “more organised” white matter than the body. However, axons do not always follow straight courses, they bend and fan, which influences the quantification of DTI metrics such as FA and MD (Nilsson et al. 2012; Ronen et al. 2014). High orientation dispersion in the corpus callosum results from fibres intertwining and diverging at the midline, as evidenced by a recent histological and DWI validation study (Mollink et al. 2017). Consistent with these observations, we observe high ODI and low FA in the body, whereas regions with low ODI have high FA. These observations hand-in-hand might suggest that FA and *v*_*ic*_ are most sensitive to myelin architecture in single-fibre regions, as corroborated by a recent combined histological and MRI validation study in rodents (Chang et al. 2017). Therefore, caution needs to be exercised when interpreting any differences in FA as related to “organisation”, since these DTI metrics do not give us an indication of organisation when the fibres are bending and fanning.

Lastly, we observe that AFD follows this low-high-low pattern in the NICAP data, and to a lesser extent in the CMIND data. Compared with the NICAP profile (*Fig 4a*), the age-matched CMIND group (*Fig 4b*) has a local minima in the genu, increase towards the body, and a slight decrease in S2 before peaking at S3. These slight profile differences might be explained by the absence of susceptibility distortion correction in CMIND. We additionally note a remarkable similarity between the NICAP AFD profile and an ex-vivo human study (Aboitiz et al. 1992) of fibre density for axons > 3 μm (*Fig 4c*). This similarity might be explained by regional differences in membrane permeability, whereby larger axon diameters can lead to alterations in the radial DWI signal with subsequent regional differences in AFD (Raffelt et al. 2012). Myelin water fraction is highest in the body of the corpus callosum, likely due to the larger fibre diameters seen in this region (Liu et al. 2010; Bjornholm et al. 2017). Therefore the bigger, more myelinated axons in the body may influence the AFD measurement in this region.

**Fig. 4.**
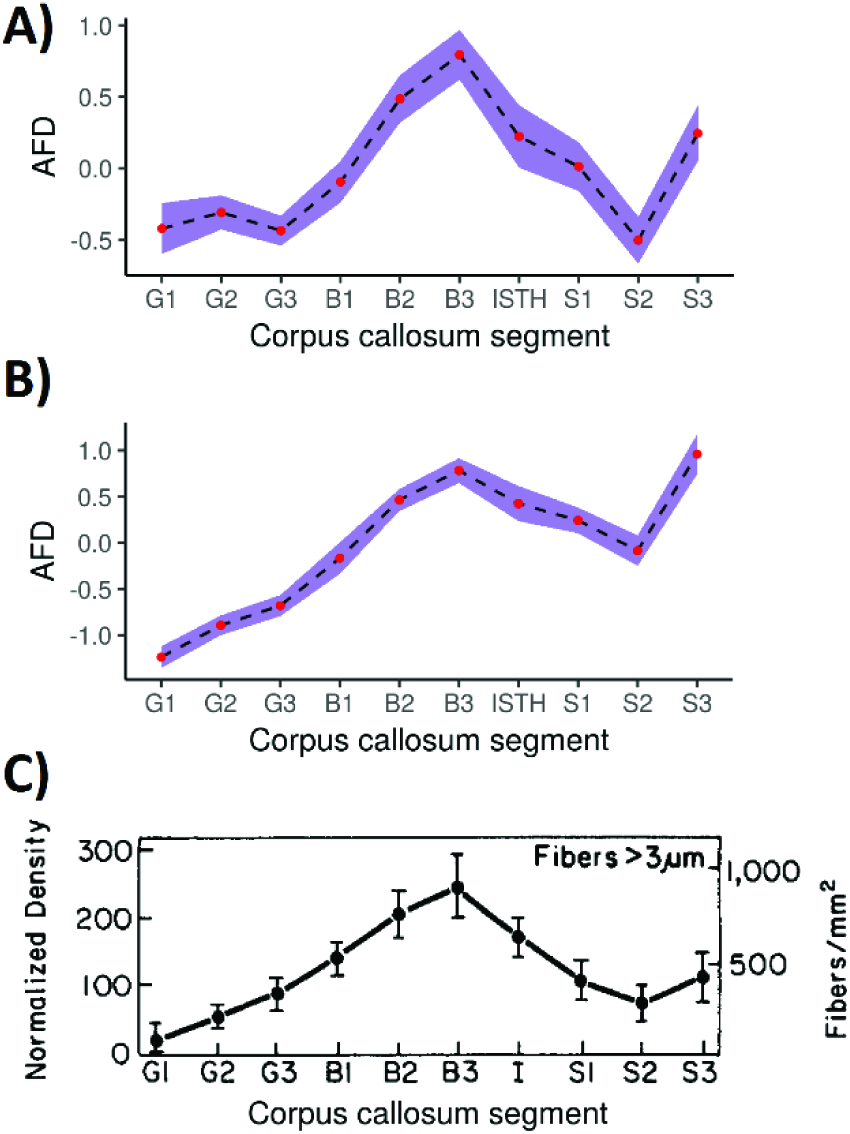
Profiles of fibre density across the corpus callosum. AFD profiles in A) NICAP dataset; and B) CMIND dataset in age-matched group. C) Aboitiz et al. (1992) light microscopy fibre density estimates for axons > 3 μm. Panel C reprinted from Aboitiz et al. (1992) with permission from Elsevier

In general, we observed a good agreement across all DTI, NODDI and AFD callosal profiles between the two cohorts, with any slight profile differences likely explained by differences in scanner acquisitions. Even in a highly organised bundle such as the corpus callosum, orientation dispersion and T2 relaxation can influence quantitative metrics, such that regions cannot be compared with one another in terms of their “organisation”. Therefore, we have chosen to focus our interpretation of these diffusion metrics in the context of its relationship with age and sex, rather than absolute comparisons between regions across the profile.

### Age-related development

Age-related development of white matter has been established in terms of volumetric (Mills et al. 2016; Lenroot and Giedd 2006), DTI (Lebel and Beaulieu 2011; Lebel et al. 2008) and NODDI (Genc et al. 2017a; Mah et al. 2017; Chang et al. 2015) studies. In general, white matter volume and microstructural organisation increases with age in the majority of white matter tracts. Our results in the CMIND cohort are in line with these previous findings, as we report significant correlations between age and each of our diffusion metrics. AFD, however, exhibited the strongest relationship with age, relative to DTI and NODDI metrics, across the corpus callosum. Although we found no main effect of age across the diffusion metrics in the NICAP cohort, this was most likely due to the relatively narrow age range (9.6 - 11.9 years) in this cohort.

Upon further investigation within each callosal segment, we observed a high correlation between AFD and age in all segments with the strongest relationship with age in the posterior midbody. The posterior midbody is known to have the highest density of large fibres, evidenced by *ex vivo* adult data (Aboitiz et al. 1992). Given that larger axons conduct action potentials faster (Horowitz et al. 2015), this strong relationship with age might reflect the need for faster interhemispheric transfer times to enable rapid communication between developing cortices of the brain during this critical period. Higher AFD with older developmental age might therefore reflect bigger axon diameter (Raffelt et al. 2017; Genc et al. 2017b), however this would need to be verified by imaging developmental populations using a high gradient system (McNab et al. 2013) or an advanced acquisition protocol (Alexander et al. 2010) to estimate axon diameter.

### Sex differences and pubertal processes

We observe higher AFD in females compared with males across the corpus callosum, most considerably in the anterior splenium and posterior genu. This is consistent with adult *ex vivo* data, whereby neurotypical females have greater fibre density in the genu and body, compared with males (Highley et al. 1999). This suggests that the observed differences in density (AFD) might be due to females having either: (1) more axons to begin with; or (2) larger diameter axons. Previous work has shown sex differences in the growth of axon and myelin in adolescents (Perrin et al. 2009), a developmental change that could be described by males having a higher proportion of large diameter fibres (Paus and Toro 2009). Therefore, it is possible that females have a higher proportion of small diameter axons.

Sex differences in brain development can be attributed to endocrine processes associated with puberty. We found that pubertal children have greater AFD in the posterior splenium compared with pre-pubertal children, consistent with our previous whole-brain analysis (Genc et al. 2017b). The pubertal period is accompanied by regulation and remodelling of white matter (Herting et al. 2017) due to rising adrenal (Maninger et al. 2009) and gonadal (Pangelinan et al. 2016) hormones (see Juraska and Willing (2017) for a recent review). We additionally demonstrated a potential step-wise relationship between pubertal group and sex, with pre-pubertal males having the lowest AFD in the splenium, and pubertal females having the highest AFD in the splenium. This additional finding suggests that females are ahead in their axonal development, likely due to the rise in adrenal and gonadal hormones one year earlier than males (Grumbach and Styne 1998). Pubertal onset is thought to be accompanied by increasing axon diameter (Genc et al. 2017b), likely driven by surges in testosterone during this critical period of neurodevelopment (Perrin et al. 2008; Pesaresi et al. 2015).

The *organisational-activational hypothesis* suggests that the brain reorganises with not only peripubertal, but perinatal exposure to testosterone (Schulz et al. 2009). In the context of our developmental data, rather than a divergence between female and male axonal development, we present evidence that they develop at the same rate. The finding of higher AFD in females over development is likely due to earlier exposure to testosterone, both perinatally and peripubertally. We do note however that our reported sex differences are mostly driven by age-related development, so it is difficult to infer sex interactions with smaller relative effect sizes. Further work is required to investigate the longitudinal development of axon and myelin properties, particularly as clinical and typical populations may have different rates of development of axon or myelin.

### Practical considerations and optimising study design

Deciding on whether to investigate myelin and/or axon depends on the question at hand. It may make more sense to quantify myelin in the case of a demyelinating disease, or axon in the case of traumatic brain injury recovery. As Jones et al. (2013) has argued, DTI is a sensitive but not specific marker of tissue microstructure. Problems only arise when making any claims about the tissue properties using DTI alone, therefore more specific markers of tissue microstructure should be included to further inform interpretations.

Practical considerations of modelling diffusion metrics depend on the acquisition parameters of the data collected, if a retrospective analysis is planned. Estimating intra-cellular volume fraction (*v*_*ic*_) obtained from NODDI requires multiple DWI shells for the compartment modelling, however ODI can be estimated using one DWI shell (Zhang et al. 2012). AFD estimation is recommended using a higher b-value to optimise signal attenuation of the extra-axonal space (e.g. b = 3000 s/mm^2^). In areas of multiple fibre orientations (basically all areas outside of the corpus callosum) the fixel-based analysis framework should be implemented to perform fibre-specific statistical comparisons (Raffelt et al. 2017; Raffelt et al. 2015). We stress that investigations should be tailored based on the acquisitions parameters of collected data - apart from the case of a prospective design whereby acquisition of multiple DWI shells can be planned in advance to quantify these parameters.

## 5. Limitations & future directions

A strength of the current study is the assessment of diffusion metrics in two independent developmental populations: a Melbourne, Australia population (NICAP dataset); and a Cincinnati, USA population (CMIND dataset). Analysing data collected from independent sites will inherently lead to mismatches in image acquisition parameters, which can affect processing and analysis. Despite differences in acquisition parameters (b-value and number of directions) and image pre-processing methods (no susceptibility distortion correction in CMIND), we observe good agreement between DTI and NODDI metrics (*Fig 2*, *Table S3*).

Whilst we have attempted to mitigate the impact of myelination on axon development by using AFD as a marker of intra-axonal volume fraction, T2 relaxation from myelin water can affect AFD measurements. We were not able to estimate myelin maps as part of the current study, however we believe this is vital for understanding how axon and myelin develop together. Therefore, future studies should attempt to quantify myelin to achieve a complementary analysis of regional growth of axon and myelin over development.

## 6. Conclusion

We present evidence of a strong relationship between apparent fibre density and age, over childhood and adolescent corpus callosum development, compared with other commonly derived diffusion metrics. This is likely driven by increasing axon diameter over development. We additionally observe that females have higher fibre density compared with males, which may be explained by differences in axon composition resulting from early surges in testosterone in both the perinatal and peripubertal period.

## 7. Compliance with Ethical Standards

All authors disclose no real or potential conflicts of interest.

All procedures performed in studies involving human participants were in accordance with the ethical standards of the institutional and/or national research committee and with the 1964 Helsinki declaration and its later amendments.

Written informed consent was obtained from the parent/guardian of all children in this study. Additionally, informed consent was obtained for adolescents that were aged 18.

## Acknowledgments

Data used in the preparation of this article were obtained from the CMIND Data Repository (Contract #s HHSN275200900018C) and NICAP study (National Health and Medical Research Council; project grant #1065895). This research and analysis was conducted within the Developmental Imaging research group, Murdoch Childrens Research Institute, supported by The Royal Children’s Hospital Foundation and the Victorian Government∙s Operational Infrastructure Support Program.

## 9. Supplementary section

**Table S1.** Multivariate profile analysis for diffusion metrics in CMIND cohort.

| Effect | $F^1$ | DF | $p$ | $\eta_p^2$ |
| --- | --- | --- | --- | --- |
| *AFD* |  |  |  |  |
| Segment | 9.85 | 9,74 | < .**.001** | .55 |
| Age | 154.14 | 1,82 | < .**.001** | .65 |
| Sex | 15.61 | 1,82 | < .**.001** | .16 |
| Age*Segment | 21.55 | 9,74 | < .**.001** | .72 |
| Sex*Segment | 0.61 | 9,74 | .78 | .07 |
| *FA* |  |  |  |  |
| Segment | 19.67 | 9,61 | < .**.001** | .74 |
| Age | 7.89 | 1,69 | .006 | .10 |
| Sex | 0.51 | 1,69 | .48 | .01 |
| Age*Segment | 2.89 | 9,61 | .03 | .25 |
| Sex*Segment | 0.99 | 9,61 | .46 | .13 |
| *MD* |  |  |  |  |
| Segment | 9.52 | 9,61 | < .**.001** | .58 |
| Age | 29.06 | 1,69 | < .**.001** | .30 |
| Sex | 1.03 | 1,69 | .31 | .02 |
| Age*Segment | 2.87 | 9,61 | .007 | .30 |
| Sex*Segment | 0.45 | 9,61 | .90 | .06 |
| *$V_{ic}$* |  |  |  |  |
| Segment | 6.62 | 9,61 | < .**.001** | .49 |
| Age | 25.18 | 1,69 | < .**.001** | .27 |
| Sex | 1.37 | 1,69 | .25 | .02 |
| Age*Segment | 0.85 | 9,61 | .57 | .11 |
| Sex*Segment | 0.88 | 9,61 | .55 | .12 |
| *ODI* |  |  |  |  |
| Segment | 21.52 | 9,61 | < .**.001** | .76 |
| Age | 11.89 | 1,69 | **.001** | .15 |
| Sex | 0.12 | 1,69 | .73 | .01 |
| Age*Segment | 4.94 | 9,61 | < .**.001** | .42 |
| Sex*Segment | 0.70 | 9,61 | .71 | .09 |

<sup>1</sup> Computed from Pillai's trace. Bold values denote $p < .005$

**Table S2.** Multivariate profile analysis for diffusion metrics in NICAP cohort.

| Effect | $F^1$ | DF | $p$ | $\eta_p^2$ |
| --- | --- | --- | --- | --- |
| *AFD* |  |  |  |  |
| Segment | 0.61 | 9,63 | .78 | .08 |
| Age | 1.22 | 1,71 | .27 | .02 |
| Sex | 0.35 | 1,71 | .56 | .01 |
| Age*Segment | 0.76 | 9,63 | .66 | .10 |
| Sex*Segment | 1.87 | 9,63 | .07 | .21 |
| *FA* |  |  |  |  |
| Segment | 1.44 | 9,63 | .19 | .17 |
| Age | 1.26 | 1,71 | .27 | .02 |
| Sex | 0.36 | 1,71 | .55 | .01 |
| Age*Segment | 1.28 | 9,63 | .27 | .15 |
| Sex*Segment | 1.97 | 9,63 | .06 | .22 |
| *MD* |  |  |  |  |
| Segment | 0.73 | 9,63 | .68 | .09 |
| Age | 0.03 | 1,71 | .86 | .01 |
| Sex | 0.33 | 1,71 | .57 | .01 |
| Age*Segment | 0.07 | 9,63 | .87 | .07 |
| Sex*Segment | 0.23 | 9,63 | .04 | .23 |
| *V<sub>ic</sub>* |  |  |  |  |
| Segment | 0.40 | 9,57 | .93 | .06 |
| Age | 0.21 | 1,65 | .65 | .01 |
| Sex | 0.13 | 1,65 | .72 | .01 |
| Age*Segment | 0.06 | 9,57 | .95 | .06 |
| Sex*Segment | 0.07 | 9,57 | .95 | .07 |
| *ODI* |  |  |  |  |
| Segment | 2.12 | 9,60 | .04 | .24 |
| Age | 0.19 | 1,68 | .66 | .01 |
| Sex | 2.59 | 1,68 | .11 | .04 |
| Age*Segment | 1.86 | 9,60 | .08 | .22 |
| Sex*Segment | 1.35 | 9,60 | .23 | .17 |

<sup>1</sup> Computed from Pillai's trace.

**Table S3.** Mean values for each diffusion metric in age-matched NICAP and CMIND groups, reported as mean (standard deviation).

| NICAP (aged 9.6 - 11.9 years, n = 74) |  |  |  |  |  |  |
| --- | --- | --- | --- | --- | --- | --- |
| Segment | AFD | FA | MD | $v_{ic}$ | ODI | |
| | $M (SD)$ | $M (SD)$ | $M (SD)$ | $M (SD)$ | $M (SD)$ | |
| G1 | .83 (.08) | .78 (.04) | .77 (.06) | .70 (.06) | .06 (.01) |  |
| G2 | .84 (.06) | .74 (.05) | .75 (.05) | .65 (.07) | .08 (.02) |  |
| G3 | .82 (.06) | .67 (.05) | .76 (.06) | .63 (.06) | .11 (.02) |  |
| B1 | .86 (.08) | .66 (.05) | .75 (.06) | .63 (.07) | .11 (.02) |  |
| B2 | .91 (.08) | .69 (.06) | .73 (.07) | .64 (.07) | .11 (.02) |  |
| B3 | .94 (.09) | .70 (.06) | .75 (.07) | .64 (.07) | .10 (.01) |  |
| ISTH | .89 (.10) | .65 (.08) | .85 (.10) | .60 (.09) | .09 (.02) |  |
| S1 | .87 (.10) | .77 (.06) | .77 (.06) | .65 (.07) | .06 (.02) |  |
| S2 | .82 (.09) | .82 (.04) | .71 (.06) | .71 (.08) | .05 (.01) |  |
| S3 | .89 (.09) | .79 (.05) | .73 (.06) | .75 (.07) | .07 (.02) |  |
| CMIND (aged 9.5 - 12.0 years, n = 15) |  |  |  |  |  |  |
| G1 | .63 (.04) | .74 (.02) | .80 (.04) | .68 (.03) | .09 (.01) |  |
| G2 | .67 (.04) | .65 (.05) | .82 (.07) | .63 (.06) | .12 (.01) |  |
| G3 | .69 (.05) | .61 (.05) | .76 (.05) | .58 (.08) | .15 (.01) |  |
| B1 | .75 (.05) | .62 (.06) | .76 (.06) | .62 (.11) | .15 (.02) |  |
| B2 | .82 (.05) | .67 (.06) | .71 (.07) | .62 (.09) | .14 (.01) |  |
| B3 | .86 (.06) | .61 (.05) | .83 (.07) | .60 (.09) | .12 (.01) |  |
| ISTH | .82 (.06) | .55 (.05) | .94 (.09) | .58 (.08) | .12 (.02) |  |
| S1 | .80 (.06) | .68 (.04) | .86 (.05) | .65 (.08) | .09 (.02) |  |
| S2 | .76 (.06) | .73 (.05) | .83 (.05) | .77 (.09) | .08 (.03) |  |
| S3 | .88 (.06) | .74 (.04) | .81 (.05) | .81 (.09) | .09 (.02) |  |

AFD values are in arbitrary units (a.u.); MD values are in $10^{-3} \text{ mm}^2/\text{s}$ .

**Table S4.** Model parameters for the linear, quadratic, and cubic fit between AFD and age in CMIND.

| Model | Linear |  | Quadratic |  | Cubic |  |
| --- | --- | --- | --- | --- | --- | --- |
| Region | *AIC* | *Adjusted R<sup>2</sup>* | *AIC* | *Adjusted R<sup>2</sup>* | *AIC* | *Adjusted R<sup>2</sup>* |
| G1 | 106 | .24 | 105.6 | .24 | 106.4 | .23 |
| G2 | 87 | .38 | 87.14 | .38 | 89.5 | .36 |
| G3 | 91.6 | .47 | 91.5 | .47 | 94.9 | .45 |
| B1 | 103.1 | .42 | 105.6 | .40 | 110 | .37 |
| B2 | 129.3 | .60 | 137.4 | .55 | 147.6 | .50 |
| B3 | 160 | .71 | 178 | .65 | 196 | .56 |
| ISTH | 167.2 | .54 | 176.1 | .48 | 185.1 | .43 |
| S1 | 146.7 | .57 | 153.1 | .53 | 161.1 | .49 |
| S2 | 156.5 | .49 | 158.7 | .47 | 163.6 | .44 |
| S3 | 188.5 | .51 | 195.5 | .47 | 203.5 | .42 |

AIC: Akaike Information Criterion; Adjusted R2: adjusted coefficient of determination for overall model.

**Fig S1.**
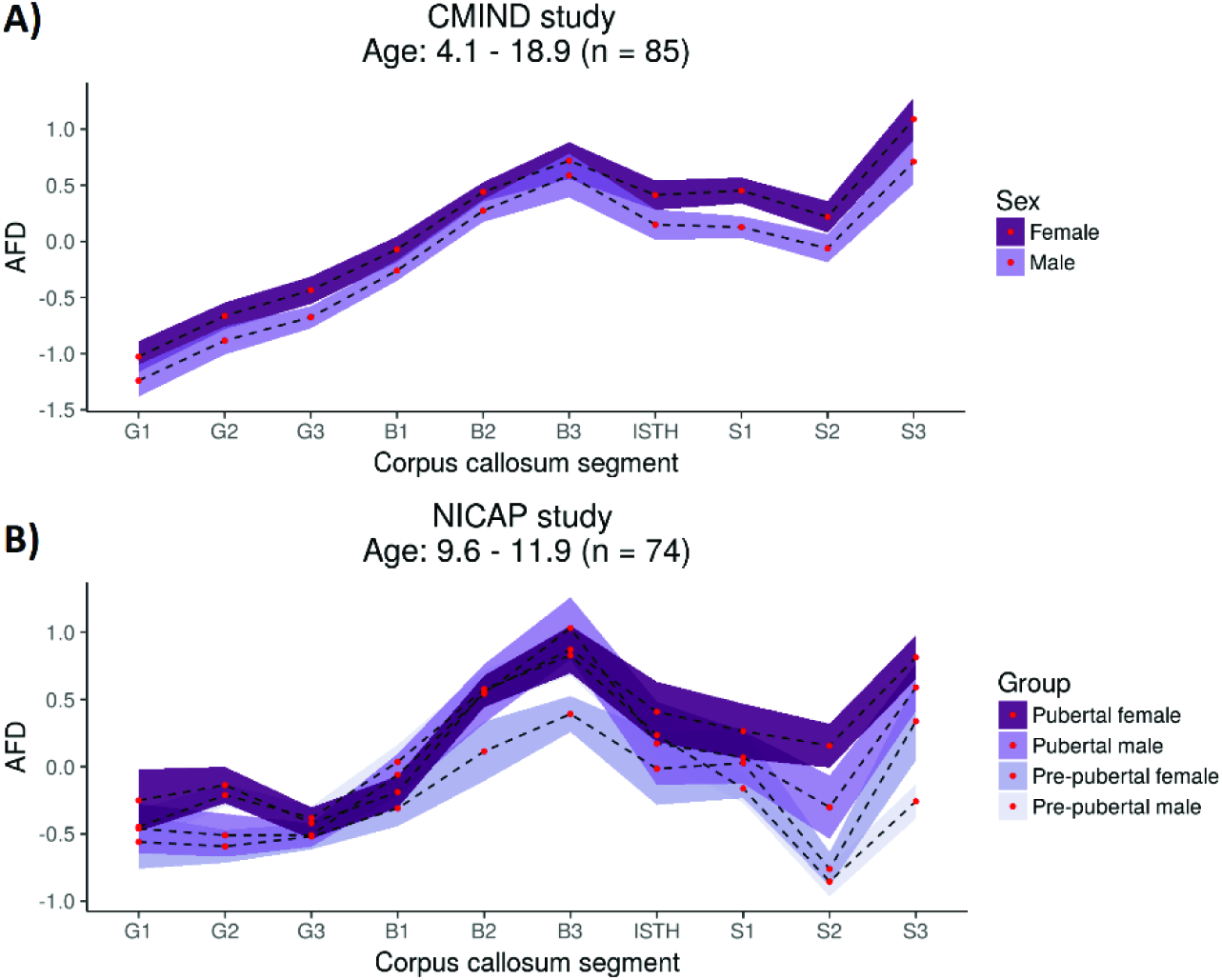
Sex and puberty differences in AFD over the corpus callosum. A) Sex differences over childhood and adolescence in CMIND, where coloured ribbons are 95% CIs. B) Pubertal type differences in, where coloured ribbons are standard error of the mean

